# The Vpu-interacting protein ATP6V0C regulates expression of tetherin and HIV-1 release

**DOI:** 10.1101/2020.03.01.972125

**Authors:** Abdul A. Waheed, Maya Swiderski, Ali Khan, Ariana Gitzen, Ahlam Majadly, Eric O. Freed

## Abstract

The HIV-1 accessory protein Vpu enhances virus release by down-regulating cell surface expression of the host restriction factor tetherin. To further understand the role of host proteins in Vpu function, we carried out yeast two-hybrid screening and identified the V0 subunit C of vacuolar ATPase (ATP6V0C) as a Vpu-binding protein. To examine the role of ATP6V0C in Vpu-mediated tetherin degradation and HIV-1 release, we knocked down ATP6V0C expression in HeLa cells and observed that ATP6V0C depletion impairs Vpu-mediated tetherin degradation, resulting in a defect in HIV-1 release. We also observed that overexpression of ATP6V0C stabilizes tetherin expression. This stabilization is specific to ATP6V0C, as overexpression of another subunit of the vacuolar ATPase, ATP6V0C”, had no effect on tetherin expression. ATP6V0C overexpression did not stabilize CD4, another target of Vpu-mediated degradation. Immunofluorescence localization studies showed that the ATP6V0C-stabilized tetherin is sequestered in a CD63- and LAMP1-positive intracellular compartment. These data demonstrate that the Vpu-interacting protein ATP6V0C plays a role in regulating tetherin expression and HIV-1 assembly and release.

## Introduction

A number of host proteins possessing antiviral activity have evolved as the first line of defense to suppress the replication of viruses in a cell-autonomous manner. These host proteins, which are often referred to as restriction factors, include APOBEC and SERINC family members, TRIM5α, SAMHD1, and tetherin. Many of these antiviral factors are either expressed constitutively or are induced by type-I interferon (IFN) (in the case of the so-called IFN-stimulated genes, or ISGs) as a component of the innate immune system (1–4). Tetherin [also known as bone marrow stromal antigen-2 (BST-2), cluster of differentiation 317 (CD317) or HM1.24] is an IFN-inducible, type II transmembrane (TM) glycoprotein that interferes with the late stage of the virus replication cycle by tethering virions to the cell surface (5,6). Tetherin was originally identified as a marker for bone marrow stromal cells; it is expressed constitutively in terminally differentiated B cells and T cells, monocytes, and dendritic cells, and is upregulated in some cancer cells (7–12). Tetherin comprises 180 amino acids and is localized in lipid rafts at the cell surface and on intracellular membranes (11,13). Tetherin has an unusual topology: it contains an N-terminal cytoplasmic tail (CT), an α-helical TM domain followed by an extracellular coiled-coil (CC) domain, and a putative C-terminal glycosylphosphatidylinositol (GPI) anchor. Because tetherin has two membrane anchors (TM and GPI-anchor) it is associated with the plasma membrane, specifically in cholesterol- and sphingolipid-rich membrane microdomains (13). The CC ectodomain of human tetherin contains three Cys residues that are essential for the formation of homodimers via disulfide bonding, and two Asn residues that are modified with N-linked oligosaccharides (13–15). Both membrane anchors, and CT and CC domains, and the three Cys residues in the CC domain are essential for the antiviral activity of tetherin (5,15,16). We reported previously that high-mannose modification of a single asparagine residue is necessary and sufficient, whereas complex-type glycosylation is dispensable, for cell-surface expression and antiviral activity of tetherin (17). Tetherin inhibits the release of a variety of enveloped viruses including not only HIV-1 but also other lentiviruses, other retroviruses, and alphaviruses, filoviruses, arenaviruses, paramyxoviruses, rhabdoviruses, flaviviruses, orthomyxoviruses, orthohepadnaviruses, and herpesviruses (reviewed in (18–20)).

Several lentiviral proteins have acquired the ability to counteract the antiviral activity of tetherin. The envelope (Env) glycoprotein of HIV-2 and some strains of simian immunodeficiency virus (SIV) counteracts tetherin by sequestering it in a perinuclear compartment, thereby down-regulating its expression from the cell surface (21–23). The Nef proteins from SIVcpz and SIVgor antagonize the respective non-human primate tetherin orthologs by decreasing their cell surface expression without inducing their degradation, possibly by intracellular sequestration (24–26). The glycoprotein M of herpes simplex virus 1 (HSV-1), the Env glycoproteins of equine infectious anemia virus (EIAV), feline immunodeficiency virus (FIV), and HERV-K human endogenous retrovirus, K5 of Kaposi’s sarcoma-associated herpesvirus (KSHV), and the non-structural protein 1 (nsP1) of chikungunya virus (CHIKV) antagonize tetherin by distinct mechanisms (27–31). HIV-1 Vpu antagonizes human, chimpanzee, and gorilla tetherin but is relatively inactive against tetherin orthologs from other non-human primates and non-primate species (25,32,33). Mechanisms by which Vpu antagonizes tetherin include (i) removing it from virus budding sites, (ii) promoting its degradation and/or (iii) down-regulating its cell-surface expression.

The accessory protein Vpu is a 16-kDa, 81-amino-acid type I integral membrane phosphoprotein containing a short luminal N-terminal domain, a 23-amino acid TM domain, and a long CT (34,35). The CT of Vpu consists of two α-helices linked by a short loop that contains two serine residues (S52 and S56) that undergo phosphorylation. Vpu is primarily localized in the endoplasmic reticulum (ER), trans-Golgi network (TGN), endosomal membranes, and to some extent at the plasma membrane (36,37). Two main functions have been attributed to Vpu: (i) proteasomal degradation of newly synthesized CD4 receptor (38–40) and (ii) down-regulation of tetherin from virus assembly sites to promote the release of viral particles (5,6). The interaction of Vpu and CD4 via their CTs leads to the recruitment of beta-transducin repeats-containing protein (β-TrCP), followed by CD4 ubiquitylation and proteasomal degradation (41), whereas interaction between Vpu and tetherin is mediated by their TM domains (32). Other functions of Vpu include downregulation of a number of host cell proteins, including major histocompatibility complex class II, tetraspanin family proteins and P-selectin glycoprotein ligand-1, from the cell surface (42–45). Vpu-mediated downregulation of CD4 and tetherin from the cell surface helps to protect HIV-1-infected cells from antibody-dependent cell-mediated cytotoxicity (42–44,46).

To better understand Vpu function, and identify Vpu-interacting cellular proteins, we previously performed a yeast two-hybrid (Y2H) screen with HIV-1 Vpu as bait. We confirmed that small glutamine-rich tetratricopeptide repeat (TPR)-containing protein α (SGTA) is a Vpu-binding protein (47,48). Overexpression of SGTA impaired HIV-1 release independent of Vpu and tetherin expression (47–50), whereas the depletion of SGTA had no significant effect on HIV-1 release or Vpu-mediated tetherin degradation (48). Further, we reported that overexpression of SGTA in the presence of Vpu induced a marked stabilization and cytosolic accumulation of the non-glycosylated form of tetherin. This accumulation of non-glycosylated tetherin in the presence of SGTA was due to inhibition of its degradation, partly due to a block in the translocation of tetherin across the ER membrane resulting in tetherin accumulation in the cytosol (48). The C-terminus of SGTA, the membrane-proximal basic residues of Vpu, and the transmembrane domain of human tetherin are required for SGTA-mediated stabilization of non-glycosylated tetherin (48).

In this study, we report the V0 subunit C of vacuolar ATPase (ATP6V0C) as another Vpu-interacting protein identified in our Y2H screen. The vacuolar H^+^-ATPase (V-ATPase) is a large, membrane-associated, multisubunit enzyme complex that functions as an ATP-dependent proton pump that acidifies intracellular compartments such as endosomes, lysosomes, Golgi-derived vesicles, clathrin-coated vesicles, synaptic vesicles, multivesicular bodies, and secretory vesicles (reviewed in (51–53)). Acidification of vacuolar compartments plays an important role in several cellular processes such as protein sorting, endocytosis, and macromolecular processing and degradation. The V-ATPase is composed of two domains: a cytoplasmic V1 domain and a transmembrane V0 domain. The V1 domain, which is responsible for ATP hydrolysis, is composed of eight subunits (A-H, in a stoichiometry of A_3_B_3_CDEFG_2_H_1-2_); the V0 subunit, which forms the H^+^ transporting channel, is composed of five subunits (a, b, c, c’, c” in a stoichiometry of adc_4_c’c” (54–56). ATP hydrolysis catalyzed by subunit V1 drives rotation of the D-F axle, which induces proton translocation across the membrane by rotating the ring of c subunits. ATP6V0C, a 16-kDa protein with four TM domains (53), has been reported to form gap junctions (57,58), and interacts with a number of cellular and viral proteins independent of other V-ATPase subunits; these include β-integrin (59), and the bovine papillomavirus E5 oncoprotein (60,61). Knockdown of ATP6V0C induced a synergistic growth-inhibitory effect in the presence of a candidate anti-cancer therapeutic in colorectal cancer cells (62), and attenuated proliferation, invasion, and glucose metabolism in esophageal cancer cells (63). Bafilomycin A1, an inhibitor of V-ATPase activity, binds ATP6V0C and inhibits proton translocation into the lysosomal lumen, thus inhibiting lysosomal acidification and cargo degradation (64). Inhibition of endosomal acidification by bafilomycin A1 abolished the infection of several viruses such as dengue virus (65), Zika virus (66), human papillomavirus (67), rhinovirus (68), equine infectious anemia virus (69), murine leukemia virus (70), and influenza A virus (71–73).

In this study, we investigated the role of ATP6V0C in Vpu-mediated tetherin degradation and HIV-1 release. Because ATP6V0C is involved in lysosomal degradation, and Vpu promotes the lysosomal degradation of tetherin, we investigated the effect of ATP6V0C depletion on HIV-1 particle production in the presence and absence of Vpu. We observed that knockdown of ATP6V0C in HeLa cells impairs Vpu-mediated tetherin degradation, thereby inhibiting HIV-1 release. We also observed that overexpression of ATP6V0C resulted in the stabilization of tetherin expression by inducing its sequestration in CD63- and LAMP1-enriched intracellular compartments. Our results demonstrate that the Vpu-interacting protein ATP6V0C plays a role in Vpu-mediated tetherin degradation and HIV-1 release.

## Results

### Identification of ATP6V0C as a Vpu-interacting protein

To identify cellular factors that are involved in Vpu function, we carried out Y2H screening using full-length Vpu as bait (for details, see experimental procedures). A number of putative Vpu-interacting proteins were identified. Previously, we reported the role of the Vpu-interacting protein SGTA in Vpu-mediated tetherin degradation and HIV-1 release (48). In this study, we investigated the role of another Vpu-interacting protein, ATP6V0C, in Vpu-mediated tetherin degradation and HIV-1 release. First, to confirm the interaction between Vpu and ATP6V0C identified by Y2H screening, we carried out co-immunoprecipitation assays. HEK293T cells were transfected with a FLAG-tagged ATP6V0C expression vector with or without a Vpu expression vector, and cell lysates were immunoprecipitated with anti-FLAG antibodies followed by immunoblotting with anti-FLAG and anti-Vpu antibodies. As shown in Fig. 1A, consistent with the Y2H result, Vpu co-immunoprecipitated with ATP6V0C (lane 4, upper right panel). To further confirm this interaction, we performed reciprocal co-immunoprecipitation assays. HEK293T cells were transfected with the Vpu expression vector in the presence and absence of ATP6V0C-expressing plasmid, and cell lysates were immunoprecipitated with anti-Vpu antibodies followed by immunoblotting with anti-Vpu and anti-FLAG antibodies. Again, ATP6V0C co-immunoprecipitated with Vpu (Fig. 1B, lane 4, lower right panel). These co-immunoprecipitation data confirm the Y2H result and demonstrate the interaction between Vpu and ATP6V0C.

**Fig. 1.**
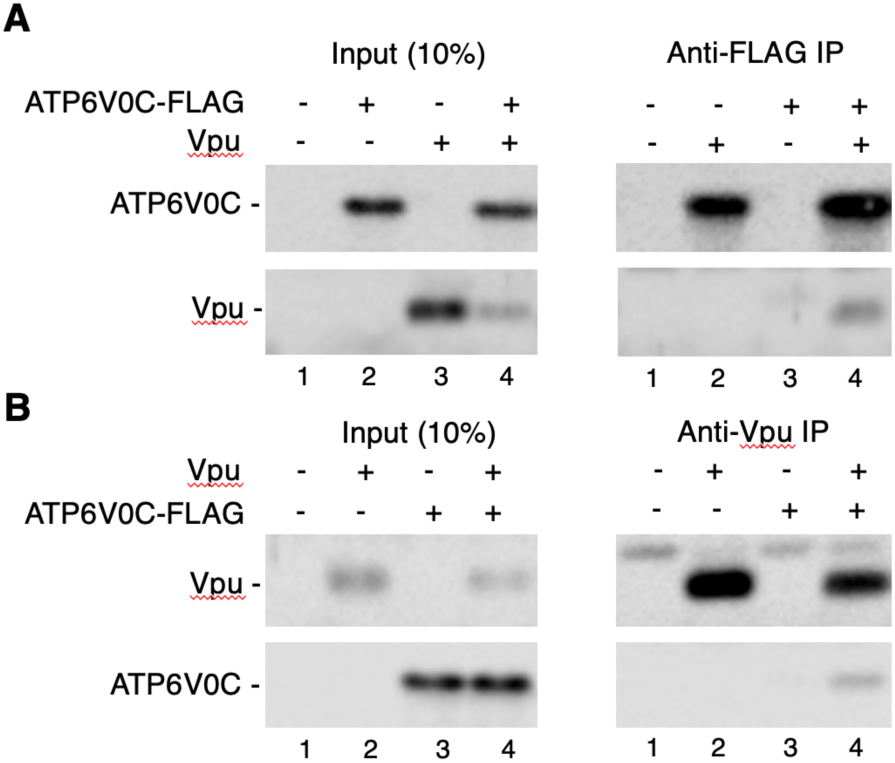
ATP6V0C co-immunoprecipitates with Vpu. 293T cells were transfected with vectors expressing FLAG-tagged ATP6V0C or Vpu alone or in combination. Twenty-four h post-transfection, cell lysates were prepared and immunoprecipitated with agarose beads conjugated with anti-FLAG (**A**) or anti-Vpu (**B**) antibodies. Complexes were washed and both cell lysates and immunoprecipitated samples were subjected to immunoblotting with anti-FLAG and anti-Vpu antibodies to detect FLAG-tagged ATP6V0C and Vpu, respectively.

### Overexpression of ATP6V0C has no effect on HIV-1 release

In our previous study, we observed that overexpression of the Vpu-interacting protein SGTA inhibited HIV-1 release in a Vpu- and tetherin-independent manner (48). To investigate whether overexpression of ATP6V0C also inhibits HIV-1 particle release we co-transfected HEK293T cells with a vector expressing FLAG-tagged ATP6V0C in the absence and presence of a tetherin expression vector and WT or Vpu-deleted (delVpu) HIV-1 molecular clones. One day posttransfection, cell and viral lysates were subjected to western blot analysis. Overexpression of ATP6V0C had no significant effect on the release of either WT or delVpu HIV-1 (Fig. 2A and B, lanes 1-3 and 7-9) in the absence of tetherin expression. In the presence of tetherin, as expected, the release of delVpu HIV-1 was reduced, but co-expression of ATP6V0C had little or no additional effect on the release of either WT or delVpu HIV-1 (Fig. 2A and B, lanes 4-6 and 10-12). However, we observed that ATP6V0C overexpression in the context of delVpu HIV-1 led to a marked increase in the levels of tetherin expression, in particular a 26-kDa tetherin species (Fig. 2A, lanes 11, and 12). We also observed an increase in tetherin expression in the presence of WT HIV-1 (Fig. 2A, lanes 5 and 6). As expected, in the absence of Vpu (Fig. 2A, lanes 10-12), higher molecular weight tetherin species were observed due to lack of Vpu-mediated tetherin degradation. These results demonstrate that ATP6V0C overexpression markedly increases tetherin levels.

**Fig. 2.**
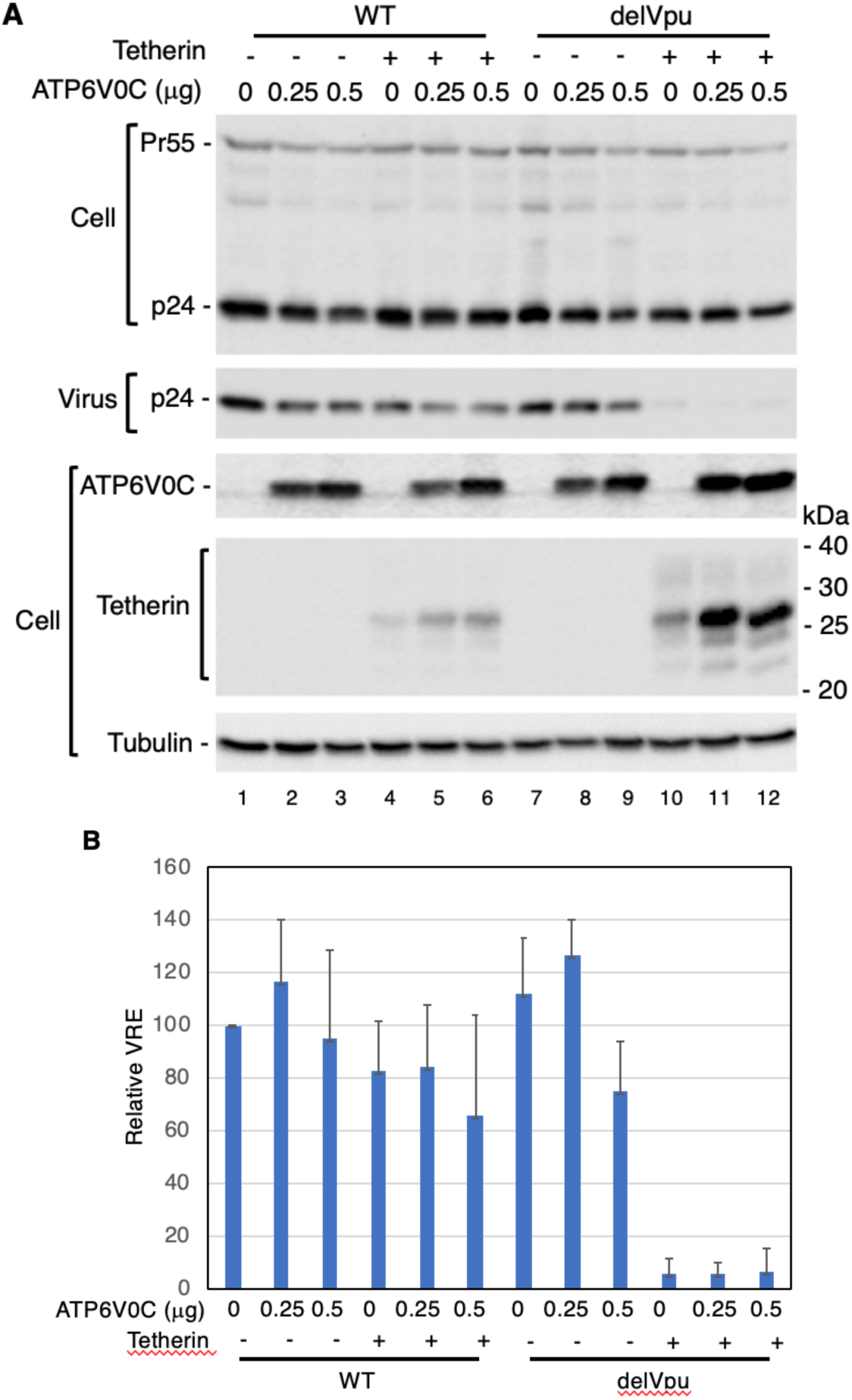
ATP6V0C overexpression has no effect on HIV-1 release. **(A)** 293T cells were transfected with WT or Vpu-defective (delVpu) pNL4-3 HIV-1 molecular clones with or without vectors expressing FLAG-tagged ATP6V0C and HA-tagged human tetherin. One day posttransfection, cell and viral lysates were prepared and subjected to western blot analysis with HIV-Ig to detect the Gag precursor Pr55Gag (Pr55) and the CA protein p24, or anti-FLAG to detect FLAG-tagged ATP6V0C, anti-HA to detect HA-tagged tetherin, or antisera against Vpu or tubulin. Mobility of molecular mass standards is shown on the right of the anti-HA blot. (**B**) Virus release efficiency (VRE) was calculated as the amount of virion-associated p24 (CA) relative to total Gag in cell and virus. VRE was set to 100% for WT pNL4-3 in the absence of ATP6V0C and tetherin. Data shown are ± SD from three independent experiments.

### Overexpression of ATP6V0C increases the levels of tetherin but reduces CD4 expression

In the above experiment, ATP6V0C and tetherin expression vectors were co-transfected with the full-length HIV-1 molecular clone. Next, we tested the expression of tetherin in the presence of ATP6V0C and Vpu in the absence of other HIV-1 proteins. HEK293T cells were transfected with tetherin expression vectors in the presence and absence of plasmids expressing ATP6V0C and Vpu. As shown in Fig. 3A, overexpressing ATP6V0C markedly increased the abundance of tetherin both in the absence and presence of Vpu. Next, we tested the effect of ATP6V0C overexpression on levels of CD4, a protein that, like tetherin, is downregulated by Vpu (40). Unlike tetherin, the expression of CD4 was reduced in the presence of ATP6V0C in a dose-dependent manner. As expected, co-expression of Vpu reduced the levels of CD4, and co-expression of ATP6V0C and Vpu further reduced CD4 levels. These results suggest that two Vpu-downregulated proteins, tetherin and CD4, are regulated differentially by ATP6V0C. As observed earlier (Fig. 1), in this experiment we also observed that Vpu levels were markedly reduced by ATP6V0C overexpression (Fig. 3B). However, the reduction in Vpu levels induced by ATP6V0C overexpression was not seen when tetherin was co-expressed (as seen in Fig. 3A). These results suggest that ATP6V0C down-regulates Vpu expression but co-expression of tetherin prevents Vpu degradation.

**Fig. 3.**
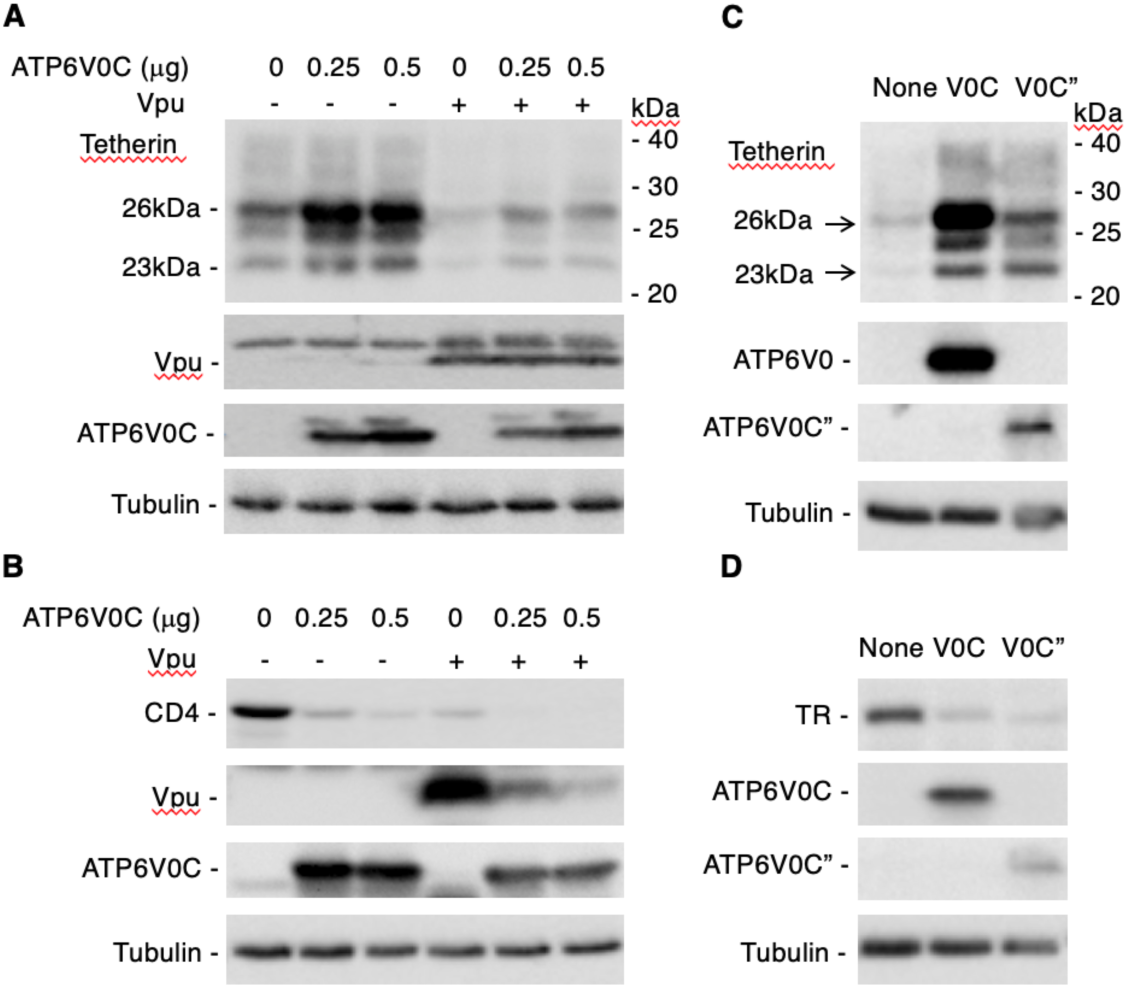
Overexpression of ATP6V0C increases the levels of tetherin but reduces CD4 and TR expression. 293T cells were transfected with vectors expressing (**A**) HA-tagged tetherin without or with FLAG-tagged ATP6V0C and Vpu expression vectors; (**B**) CD4 without or with FLAG-tagged ATP6V0C and Vpu expression vectors; (**C**) HA-tagged tetherin without or with FLAG-tagged ATP6V0C or ATP6V0C” expression vectors; (**D**) transferrin receptor (TR) without or with FLAG-tagged ATP6V0C or ATP6V0C” expression vectors. Twenty-four h post-transfection, cell lysates were prepared and subjected to western blot analysis with anti-FLAG to detect FLAG-tagged ATP6V0C, anti-HA to detect HA-tagged tetherin, or antisera against Vpu, CD4, or TR. Molecular mass markers are shown on the right of the anti-HA blots.

We next investigated whether the increase in tetherin expression is specific for the V0C subunit of V-ATPase. To test this, we overexpressed ATP6V0C”, another V0 subunit of V-ATPase. In contrast to our observations with ATP6V0C, overexpressing ATP6V0C” modestly increased tetherin expression (3.3-fold increase with ATP6V0C” compared to 10.7-fold increase with ATP6V0C) (Fig. 3C). Because V-ATPases are involved in lysosomal degradation, we also examined the effect of overexpressing the V0C and V0C” subunits on the levels of transferrin receptor (TR), which, like tetherin, is a lysosomally degraded protein (74). In contrast to our observations with tetherin, we observed that overexpressing either V0C or V0C” subunits reduced the levels of TR (Fig. 3D). These results suggest that overexpression of V0 subunits of V-ATPase does not globally impair the lysosomal degradation pathway, but rather increases the lysosomal degradation of TR.

### Tetherin co-immunoprecipitates with ATP6V0C and this interaction is independent of Vpu

The data presented in Figs. 2A, 3A, and 3C demonstrate that overexpression of the Vpu-interacting protein ATP6V0C increases tetherin expression. Because this phenomenon takes place even in the absence of Vpu, we speculated that ATP6V0C may directly interact with tetherin. To test this hypothesis, we carried out co-immunoprecipitation assays. HEK293T cells were transfected with FLAG-tagged ATP6V0C expression vectors in the absence and presence of Vpu and HA-tagged tetherin, and immunoprecipitated with anti-FLAG antibodies followed by immunoblotting with anti-FLAG, HA and Vpu antibodies. As shown in Fig 4A, tetherin co-immunoprecipitated with ATP6V0C. As shown in Fig. 1, we also observed Vpu in the co-immunoprecipitated fraction (Fig. 4A). To confirm these results, we carried out reciprocal co-immunoprecipitation assays. HEK 293T cells were transfected with HA-tagged tetherin in the absence and presence of Vpu and FLAG-tagged ATP6V0C and immunoprecipitated with anti-HA antibodies followed by immunoblotting with anti-HA, FLAG, and Vpu antibodies. Here again, we could efficiently pull down ATP6V0C both in the absence and presence of Vpu (Fig. 4B). We also observed co-immunoprecipitation of Vpu with tetherin as reported in the literature (75,76). These results suggest that these three proteins interact independently with one another, i.e., Vpu interacts with ATP6V0C (Figs. 1 and 4), ATP6V0C interacts with tetherin (Fig.4), and tetherin interacts with Vpu (Fig. 4).

**Fig. 4.**
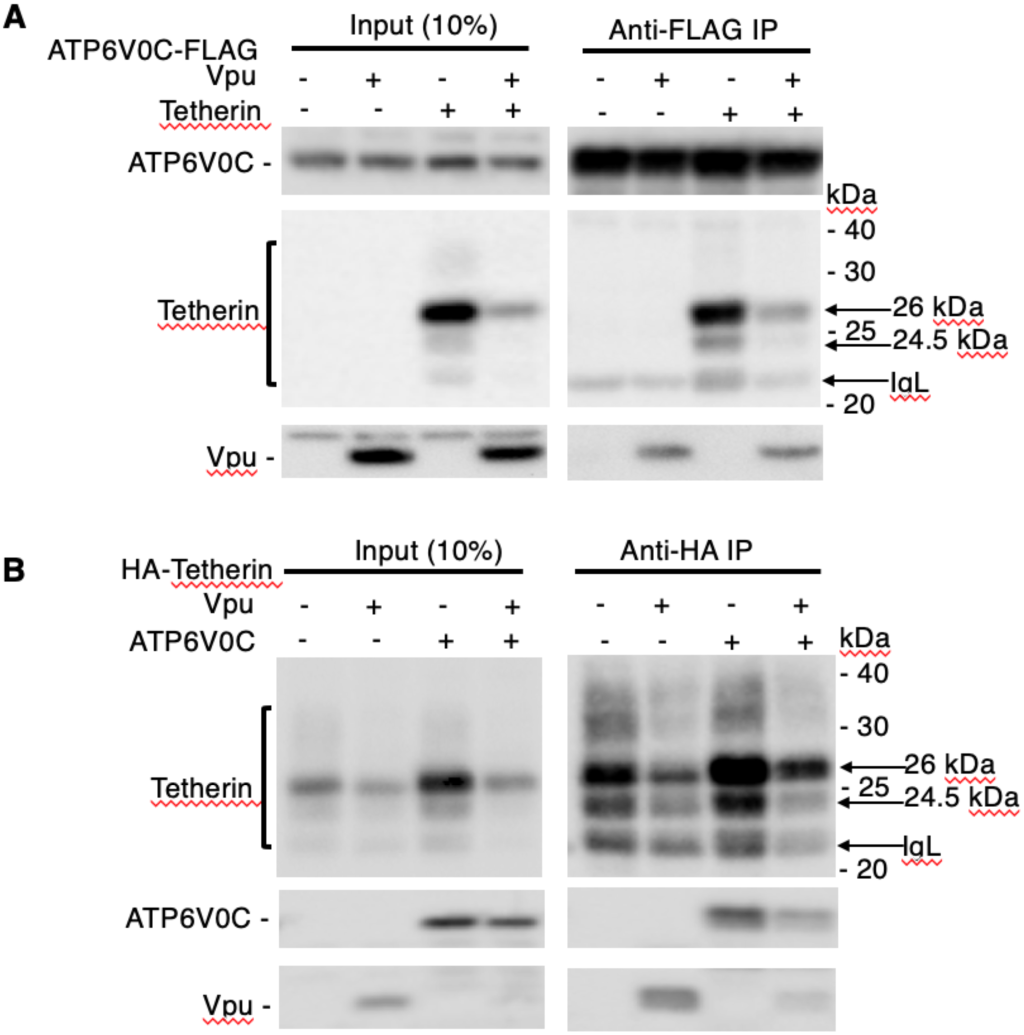
Tetherin co-immunoprecipitates with ATP6V0C independent of Vpu. 293T cells were transfected with vectors expressing FLAG-tagged ATP6V0C without or with HA-tagged tetherin and Vpu expression vectors (**A**) or HA-tagged tetherin without or with FLAG-tagged ATP6V0C and Vpu expression vectors (**B**). Twenty-four h post-transfection, cell lysates were prepared and immunoprecipitated with agarose beads conjugated with anti-FLAG (**A**) or anti-HA (**B**) antibodies. Complexes were washed and both cell lysates and immunoprecipitated samples were subjected to immunoblotting with anti-FLAG, anti-HA and anti-Vpu antibodies to detect FLAG-tagged ATP6V0C, HA-tagged tetherin, and Vpu, respectively. Mobility of molecular mass standards is shown on the right of the anti-HA blot. The location of different species of tetherin and Ig light chain (IgL) is indicated by the arrows.

### ATP6V0C overexpression stabilizes tetherin

To elucidate the mechanism by which ATP6V0C increases the levels of tetherin, we performed pulse-chase analysis. 293T cells expressing tetherin alone or coexpressing tetherin with ATP6V0C were pulse-labeled with [^35^S]Met/Cys for 30 min and chased for 0, 0.5, 1, 2, 4 h in unlabeled medium. We observed that ATP6V0C overexpression increased the half-life of tetherin (Fig. 5). Quantification indicated that the half-life of the 23- and 26-kDa tetherin species was 32 and 120 min in the absence and presence of ATP6V0C overexpression, respectively. These results indicate that ATP6V0C overexpression prevents the degradation of tetherin, leading to its stabilization and time-dependent accumulation.

**Fig. 5.**
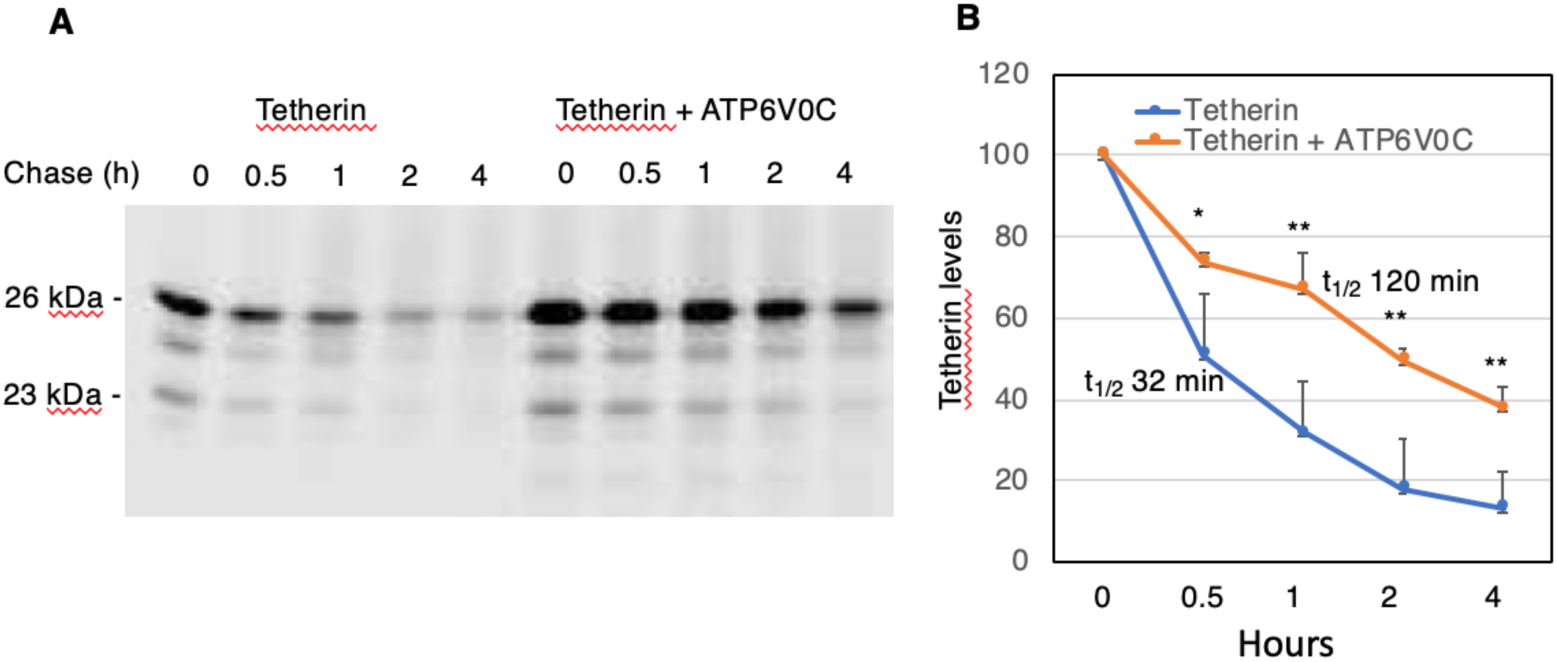
ATP6V0C overexpression stabilizes tetherin. (**A**) 293T cells were transfected with vectors expressing HA-tagged tetherin without or with FLAG-tagged ATP6V0C expression vector. 24 h post-transfection, cells were labeled with [^35^S]Met/Cys for 30 min and then chased for 0, 0.5, 1, 2, or 4 h in unlabeled medium. Cell lysates were prepared and immunoprecipitated with anti-HA antibodies and analyzed by SDS-PAGE followed by fluorography. Positions of 26-kDa (glycosylated) and 23-kDa (non-glycosylated) tetherin are indicated. (**B**) The levels of radiolabeled 23- and 26-kDa tetherin species were quantified at each time point from three independent experiments, ± SD. Tetherin levels at time zero was taken as 100%. The p values (two-tailed paired t-test) are calculated for each time point. *p < 0.05, **p < 0.02. The half-life (t_1/2_) of tetherin under each condition is indicated.

### The cytoplasmic tail, GPI-anchor, and dimerization of human tetherin are required, whereas glycosylation and the transmembrane domain are dispensable, for ATP6V0C-mediated tetherin stabilization

We next used several tetherin mutants to investigate which domain(s) of tetherin are essential for ATP6V0C-mediated tetherin stabilization. As shown in Fig. 6A, the expression of the non-glycosylated double mutant (N65,92A) is increased, whereas the non-dimerizing (CCC), cytoplasmic tail-deleted (delCT), and GPI-anchor-deleted (delGPI) mutants show no increased expression upon ATP6V0C overexpression (Fig. 6A). We also tested tetherin proteins from other species, African green monkey (Agm) and rhesus macaque, that are not downregulated by Vpu due lack of Vpu-tetherin interaction. Unlike human tetherin, Agm and rhesus tetherins were not stabilized. Chimeras in which the transmembrane domain of human tetherin was replaced with the corresponding sequences from Agm or rhesus tetherin (Hu-Agm and Hu-Rh, respectively) were also stabilized by ATP6V0C overexpression. These results indicate that the transmembrane domain and glycosylation of human tetherin are dispensable whereas the cytoplasmic tail, GPI-anchor, and dimerization of tetherin are required for ATP6V0C-mediated stabilization.

**Fig. 6.**
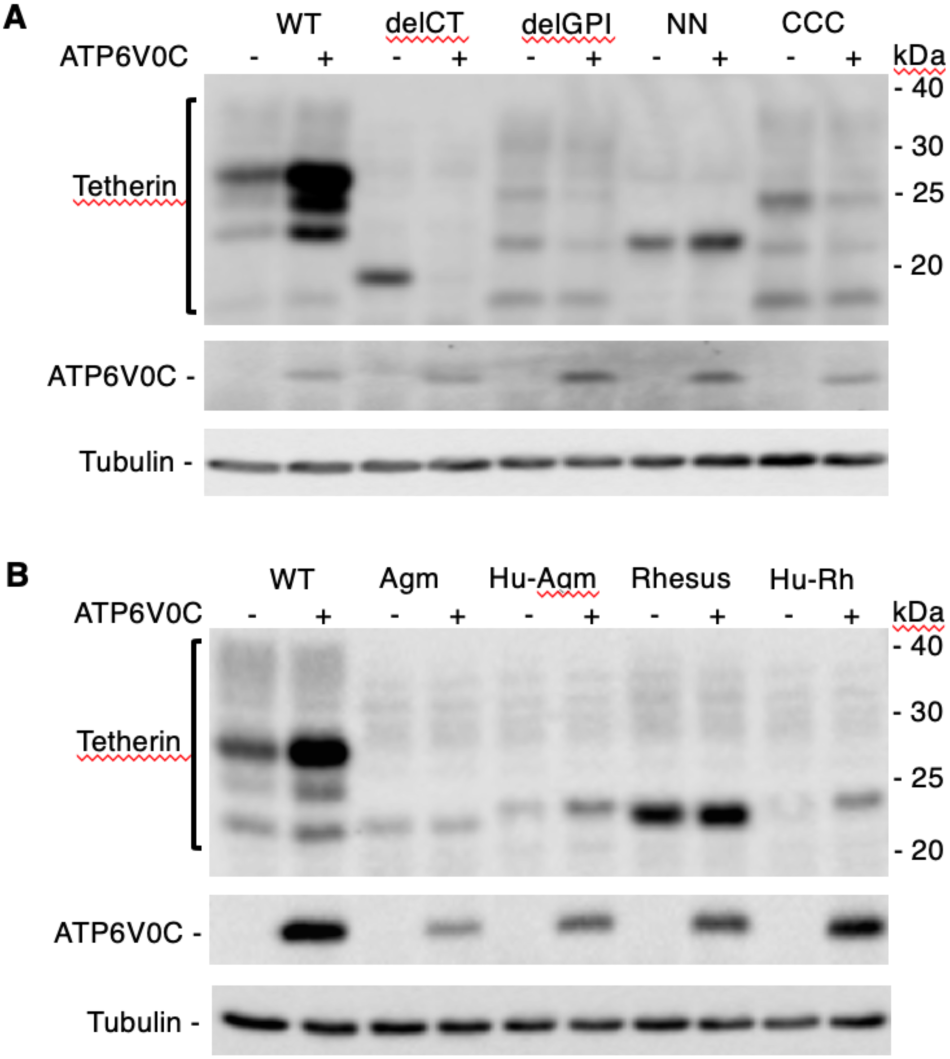
The cytoplasmic tail, GPI-anchor, and dimerization of human tetherin are required for ATP6V0C-mediated tetherin stabilization. 293T cells were transfected with vectors expressing HA-tagged tetherin with or without FLAG-tagged ATP6V0C expression vector. Tetherin variants tested include: (**A**) a cytoplasmic tail deletion mutant (delCT), a GPI-anchor deletion mutant (delGPI), non-glycosylated mutant (NN), and non-dimerizing mutant (CCC); (**B**) Agm tetherin (Agm), Rhesus tetherin (Rh), and a human tetherin containing the Agm (Hu-Agm) or Rh (Hu-Rh) transmembrane domains. One day posttransfection, cell lysates were prepared and immunoblotted with anti-HA to detect HA-tagged tetherin, anti-FLAG to detect FLAG-tagged ATP6V0C, or anti-tubulin antisera. Mobility of molecular mass standards is shown on the right of the anti-HA blot.

### Stabilization of tetherin is not due to inhibition of lysosomal degradation

The results presented above (Figs. 3A, B and D) demonstrate that although ATP6V0C overexpression stabilizes tetherin expression, it reduces the levels of two lysosomally degraded proteins, CD4 and TR. To explore further the connection between ATP6V0C overexpression and lysosomal function, we treated cells with bafilomycin, which specifically binds the ATP6V0C subunit and prevents proton transport and lysosomal acidification (77). We observed an increase in the expression of tetherin in bafilomycin-treated cells (Fig. 7A). Strikingly, the expression of tetherin is much higher in ATP6V0C overexpressing cells that are treated with bafilomycin relative to cells not treated with the lysosomal inhibitor. The higher expression of tetherin in ATP6V0C-expressing cells treated with bafilomycin could be due to the additive effect of tetherin stabilization by ATP6V0C and inhibition of degradation by bafilomycin. Next, we tested the effect of overexpressing ATP6V0C on Vpu expression. As shown in Fig. 7B, the expression of Vpu is reduced by co-expressing ATP6V0C. These results confirm our earlier observation in Fig. 3B, which was made in the presence of CD4 (Fig. 3B). Treating these cells with bafilomycin increases the expression of Vpu, suggesting that ATP6V0C overexpression leads to degradation of Vpu through the lysosomal pathway (Fig. 6B). This reduction in the expression of Vpu induced by ATP6V0C overexpression is not observed in the presence of tetherin (Fig. 2A lanes 5 and 6, and Fig. 3A, lanes 5 and 6). These observations suggest that tetherin binding to ATP6V0C reduces the ability of ATP6V0C to down-regulate Vpu, consistent with our finding that ATP6V0C, Vpu, and tetherin form a trimeric complex.

**Fig. 7.**
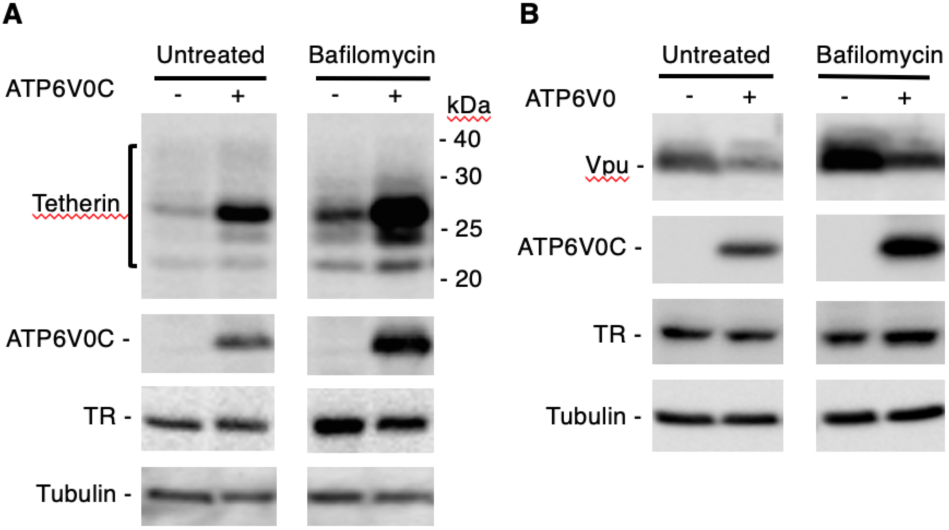
Stabilization of tetherin is not due to inhibition of lysosomal degradation. 293T cells were transfected with vectors expressing HA-tagged tetherin (**A**) or Vpu (**B**) without or with FLAG-tagged ATP6V0C expression vector. Eight h posttransfection, cells were either untreated or treated with bafilomycin for 18 h prior to cell lysis and immunoblotting with anti-HA to detect HA-tagged tetherin, anti-FLAG to detect FLAG-tagged ATP6V0C, or anti-TR or anti-tubulin antisera. Molecular mass markers are shown on the right of the anti-HA blot.

### ATP6V0C overexpression sequesters tetherin in lysosomal and CD63-positive compartments

To investigate the effect of ATP6V0C overexpression on tetherin localization, we performed immunofluorescence microscopy analysis. 293T cells were transfected with HA-tetherin in the absence and presence of C-terminally FLAG-tagged ATP6V0C, and cells were fixed and stained with anti-HA and anti-FLAG antibodies. As reported previously (13) tetherin is normally localized predominantly on the cell surface (Fig. 8A). Upon ATP6V0C overexpression, the localization of tetherin markedly shifts from the plasma membrane to intracellular vesicular compartments. Interestingly, ATP6V0C is also localized in these compartments, resulting in high colocalization of tetherin and ATP6V0C [Pearson correlation coefficient (R) value 0.81±0.09 (Fig. 8A)]. We next characterized the compartments in which tetherin is sequestered by ATP6V0C. 293T cells expressing tetherin in the absence and presence of ATP6V0C were stained for tetherin and the trans-Golgi network (TGN) marker TNG46, the late endosome marker CD63, or lysosomal marker LAMP-1. We observed little or no colocalization between tetherin and TNG46 (data not shown); however, we observed high levels of co-localization of tetherin with both CD63 [(Fig. 8B), R=0.757±0.088] and LAMP-1 [(Fig. 8C) R=0.758±0.089]. These results indicate that ATP6V0C sequesters tetherin in a compartment that is positive for late endosomal and lysosomal markers. That ATP6V0C induces accumulation of tetherin in CD63- and LAMP-1-positive compartments without inducing tetherin degradation suggests that the late endosomal and LAMP-1-positive vesicles in which tetherin accumulates represent an aberrant, non-functional lysosomal compartment.

**Fig. 8.**
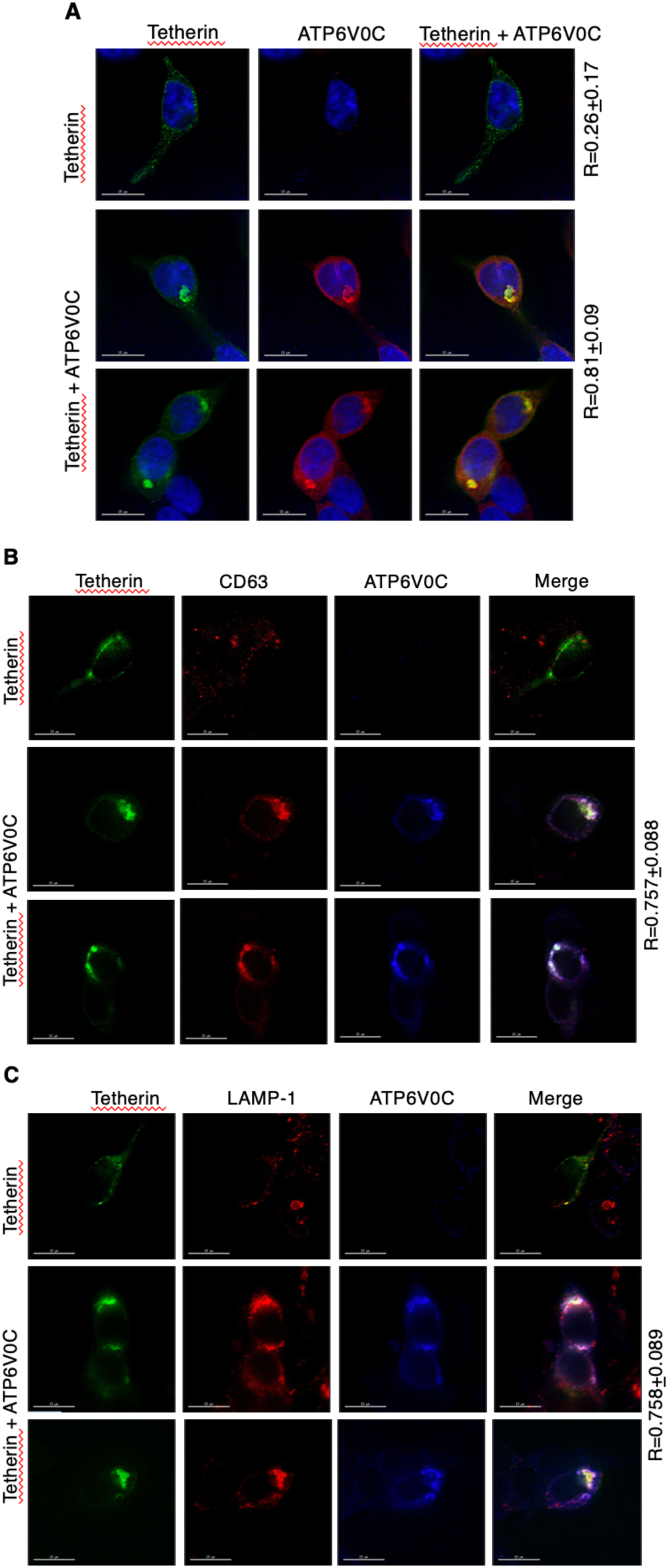
ATP6V0C overexpression sequesters tetherin in LAMP-1 and CD63-positive compartments. 293T cells were transfected with HA-tagged tetherin expression vector alone or in combination with the FLAG-tagged ATP6V0C expression vector. Twenty-four h post-transfection cells were fixed and stained with (**A**) anti-HA (green), anti-FLAG (red) and DAPI (blue); (**B**) anti-HA (green), anti-CD63 (red) and anti-FLAG (blue); (**C**) anti-HA (green), anti-LAMP-1 (red) and anti-FLAG (blue) antibodies. Images were acquired with a Delta-Vision RT deconvolution microscope. Colocalization between tetherin and ATP6V0C (**A**), tetherin and CD63 (**B**), and tetherin and LAMP-1 (**C**) was quantified by calculating the Pearson correlation coefficient (R values) ± SD from 20-30 cells. Scale bars, 15 μm.

We next overexpressed individual tetherin mutants in the presence of ATP6V0C and examined the localization of tetherin and ATP6V0C. CT-deleted tetherin and the non-dimerizing tetherin mutant, which are both localized to a perinuclear region, are not sequestered by ATP6V0C (Fig. S1A and S1B). Interestingly, in the presence of these tetherin mutants, ATP6V0C itself is diffusely localized in the cytosol. The GPI-deleted tetherin mutant, which is localized on the plasma and internal membranes, is also not sequestered by ATP6V0C (Fig. S1C) and ATP6V0C localization remains cytosolic. However, the non-glycosylated tetherin mutant is sequestered and colocalizes with ATP6V0C in an internal compartment (Fig. S1D). These results correlate with our biochemical observations in Fig. 6A; i.e. tetherin molecules that are stabilized by ATP6V0C overexpression colocalize with ATP6V0C in an internal compartment, putatively an aberrant late endosome or lysosome.

### Knockdown of ATP6V0C inhibits HIV-1 release by enhancing tetherin expression

To investigate the role of endogenous ATP6V0C in tetherin expression and HIV-1 release, we knocked-down ATP6V0C expression in HeLa cells with siRNA (knock-down efficiency >90%; Fig. 9C) and analyzed the expression of tetherin and HIV-1 release. Knocking-down ATP6V0C increased the expression of endogenous tetherin, resulting in a reduction in WT virus release efficiency (Fig. 9A and B). As expected (78,79), in the absence of ATP6V0C knock-down, the release of delVpu HIV-1 is reduced in HeLa cells because of the absence of Vpu-mediated tetherin degradation (Fig. 9A and B). Knock-down of ATP6V0C increased tetherin expression and further reduced delVpu release. Knocking down tetherin along with ATP6V0C rescued the release of both WT and delVpu HIV-1, suggesting that the impaired virus release observed upon knocking down ATP6V0C is due to higher tetherin expression. These results demonstrate that ATP6V0C is required for both normal turnover of tetherin and Vpu-mediated tetherin degradation in HeLa cells. Further, to confirm that the reduction in virus release observed upon ATP6V0C knock-down in HeLa cells is due to increased levels of tetherin, we knocked-down ATP6V0C in 293T cells, which do not express tetherin. In 293T cells, we do not see any significant reduction in virus release upon knock-down of ATP6V0C (Fig 9D and E), consistent with the hypothesis that the decrease in HIV-1 release in ATP6V0C-depleted HeLa cells is due to increased tetherin expression. Interestingly, the levels of TR are not reduced by knock-down of ATP6V0C (Fig. 9D).

**Fig. 9.**
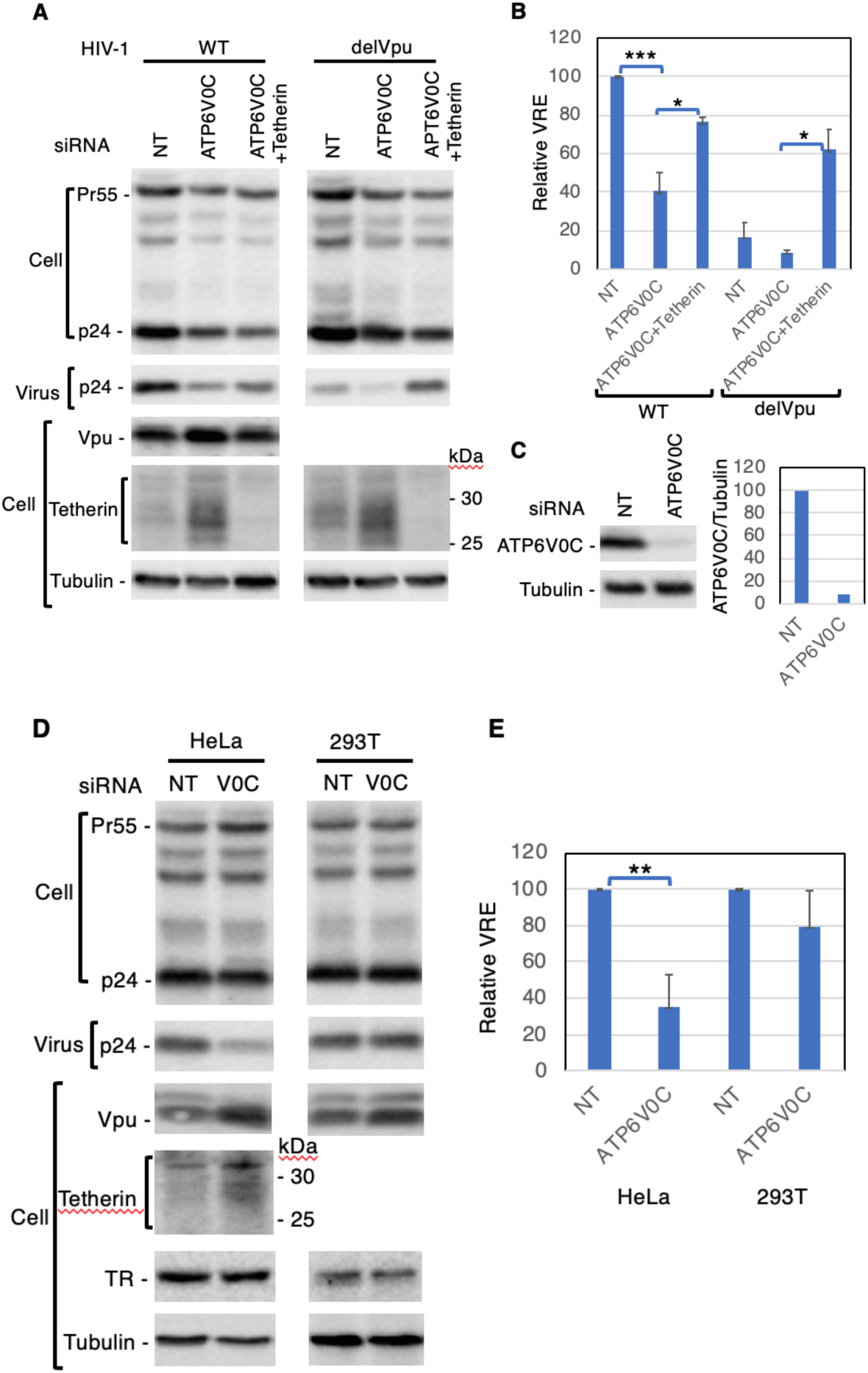
The knockdown of ATP6V0C inhibits HIV-1 release by enhancing tetherin expression. HeLa (**A and B**) or 293T (**D and E**) cells were transfected with non-targeting siRNA (NT) or ATP6V0C siRNA alone or in combination with tetherin siRNA. One-day later, cells were infected with VSV-G-pseudotyped wild-type (WT) or Vpu-defective HIV-1; after 8 h the medium was changed, and the siRNA transfection was repeated. Two days later, cell and viral lysates were prepared and subjected to western blot analysis with HIV-Ig to detect the Gag precursor Pr55Gag (Pr55) and the CA protein p24, or antibodies specific for tetherin, Vpu, or tubulin. (**B and E**), virus release efficiency (VRE) was calculated as in Fig. 2. VRE was set to 100% for WT HIV-1 in the presence of NT siRNA. (**C**) To monitor ATP6V0C knockdown efficiency, 293T cells were transfected with the FLAG-tagged ATP6V0C expression vector and treated with NT or ATP6V0C siRNA as in Fig. 9A and two-days later cells were lysed. Relative expression of FLAG-tagged ATP6V0C in the presence of NT siRNA was set to 100%. Mobility of molecular mass standards is indicated on the right of the tetherin blot. Data shown are ± SD from three to four independent experiments. P values (two-tailed paired t-test): *p < 0.02, **p < 0.01, ***p < 0.002.

## Discussion

We identified SGTA and ATP6V0C as Vpu-interacting proteins in a Y2H screen. In this study, we confirmed the interaction between Vpu and ATP6V0C using co-immunoprecipitation assays. Although in our earlier study siRNA-mediated depletion of SGTA had no significant effect on the release of either WT or Vpu-defective HIV-1 in tetherin-expressing HeLa cells, overexpression of SGTA interfered with HIV-1 release in a Vpu- and tetherin-independent manner (48). In this study, we observe that overexpression of ATP6V0C has no significant effect on HIV-1 release, whereas knockdown of ATP6V0C in HeLa cells, but not in tetherin-negative 293T cells, inhibits the release of both Vpu(+) and Vpu(-) HIV-1. The inhibition of virus release mediated by ATP6V0C depletion in HeLa cells is rescued by knockdown of endogenous tetherin, indicating that the inhibition of HIV-1 release upon knockdown of ATP6V0C is mediated by tetherin. Further, we observe that ATP6V0C depletion in HeLa cells results in elevated levels of endogenous tetherin independent of Vpu expression, indicating that endogenous ATP6V0C is not only required for Vpu-mediated tetherin degradation but also for normal tetherin turnover. We and others have reported that Vpu induces the degradation of tetherin by both proteasomal (80–83) and lysosomal (75,83,84) pathways. Because ATP6V0C regulates vesicular acidification required for lysosomal degradation (51–53), we suggest that ATPV0C is essential for Vpu-mediated lysosomal tetherin degradation to promote HIV-1 release.

We observed that ATP6V0C overexpression results in the accumulation of tetherin by preventing its degradation and that this accumulation is Vpu-independent. Overexpression of another subunit of the V-ATPase, ATP6V0C”, has only a small effect on tetherin expression compared to ATP6V0C, indicating that tetherin stabilization is specific to ATP6V0C. Further, the ATP6V0C-mediated stabilization is specific to tetherin, as overexpression of ATP6V0C did not stabilize the Vpu-downregulated protein CD4 or interfere with its Vpu-mediated down-regulation. Overexpression of ATP6V0C reduced the expression of CD4 and TR, both of which undergo lysosomal degradation (38–40). Treating ATP6V0C overexpressing cells with the lysosomal inhibitor bafilomycin, which specifically targets the ATP6V0C subunit and prevents lysosomal degradation (64), further increased tetherin expression. Overexpression of ATP6V0C reduced the expression of Vpu, which could be rescued by treating cells with bafilomycin, indicating that ATP6V0C overexpression leads to degradation of Vpu through the lysosomal pathway. These results indicate that overexpression of ATP6V0C does not exert a dominant-negative effect on lysosomal function. Our co-immunoprecipitation assays demonstrate that Vpu, ATP6V0C, and tetherin interact independently of one another. Together, these results suggest that ATP6V0C interaction with tetherin prevents its degradation, leading to its accumulation. Using deletion and point mutants, we determined that the cytoplasmic tail, GPI-anchor, and dimerization of tetherin are essential for ATP6V0C6-mediated tetherin stabilization. Immunofluorescence localization studies showed that the ATP6V0C-stabilized tetherin is sequestered in CD63- and LAMP1-enriched intracellular compartments, whereas tetherin mutants that are not stabilized by ATP6V0C are not sequestered in these internal compartments. The observation that ATP6V0C increases tetherin levels without affecting HIV-1 release is explained by the intracellular sequestration of ATP6V0C-stabilized tetherin; this stabilized tetherin is not available to inhibit particle release at the plasma membrane.

The basis for the interaction between Vpu and ATP6V0C remains unclear. Vpu could have evolved an interaction with ATP6V0C to antagonize the activity of the V-ATPase or to promote its activity in degrading Vpu-target proteins. Because ATP6V0C is a pore-forming and proton-conducting subunit of the V-ATPase, we initially anticipated that the interaction of Vpu with overexpressed ATP6V0C might facilitate Vpu-mediated tetherin degradation. However, we observed that overexpression of ATP6V0C resulted in the accumulation, rather than lysosomal degradation, of tetherin. Although overexpression of ATP6V0C increases the levels of tetherin, this effect is independent of Vpu. Interactions between virally encoded factors and ATP6V0C or other V-ATPase subunits have been reported in other systems. For example, a direct interaction between bovine and human papillomavirus protein E5 and ATP6V0C has been reported (60,85). Although several studies have demonstrated that E5 disrupts acidification of endosomes (86–88) this activity appears to be independent of E5–ATP6V0C binding (89). A human cytomegalovirus (hCMV) microRNA has been found to target ATP6V0C (90), which, paradoxically, is required for efficient hCMV virus assembly in culture (91). Why hCMV would target for degradation a host protein required for its replication has not been elucidated. The non-structural protein 3A of enterovirus 71 (EV71) interacts with ATP6V0C and this interaction is critical for EV71 replication, as knockdown of ATP6V0C or treating cells with bafilomycin inhibits propagation of EV71 (92). Dengue virus pre-membrane (prM) protein interacts with V-ATPase and this interaction is essential for both entry and egress of dengue virus (93). HIV-1 and SIV Nef have been reported to interact with the H subunit of the V-ATPase, thereby connecting Nef and the endocytosis machinery required for Nef-mediated CD4 downregulation (94–96). Finally, HTLV-1 p12I has been reported to interact with ATP6V0C (97,98); the implications of this interaction remain to be defined.

In the course of this study, we also observed that the Vpu target protein, tetherin, interacts with ATP6V0C. As a consequence of this interaction, overexpression of ATP6V0C leads to the stabilization of tetherin and the sequestration of tetherin in an internal compartment that contains CD63 and LAMP-1. Variants of tetherin that are not stabilized by ATP6V0C fail to relocalize to the internal compartment. Whereas ATP6V0C overexpression stabilizes tetherin, it causes a reduction in the levels of TR and CD4, two proteins that undergo lysosomal degradation. These results suggest that the stabilization of tetherin by ATP6V0C overexpression is a result of the (direct or indirect) interaction between tetherin and ATP6V0C and is not a consequence of global disruption of lysosomal activity.

In summary, in this report, we show that overexpressed ATP6V0C interacts with tetherin and stabilizes its expression by preventing study-state degradation and sequestration in intracellular compartments. In contrast, endogenous ATP6V0C is required for Vpu-mediated tetherin degradation and enhancement of HIV-1 release in HeLa cells. These results demonstrate that the Vpu-interacting protein ATP6V0C regulates the expression of tetherin and HIV-1 release.

## Experimental procedures

### Plasmids, antibodies, and chemicals

Vectors expressing C-terminally FLAG-tagged ATP6V0C and ATP6V0C” were obtained from OriGene Technologies Inc. (Rockville, MD). The pcDNA-Vphu vector, bearing the codon-optimized *vpu* gene, was used for expressing Vpu (99). The full-length, infectious HIV-1 molecular clone pNL4-3 and the Vpu-defective counterpart pNL4-3delVpu were described previously (35,100). pNL4-3delVpu and pcDNA-Vphu were kindly provided by K. Strebel. Vectors expressing N-terminally HA-tagged human tetherin, the deletion mutant derivatives delCT and delGPI, tetherin from African green monkey (Agm) and Rhesus (Rh), and chimeric human tetherin with transmembrane domains from Agm (Hu-Agm) or Rh (Hu-Rh) were described previously (5,32), and were generously provided by P. Bieniasz. The non-glycosylated (NN) and non-dimerizing (CCC) tetherin mutants have been described previously (48). The CD4 expression vector pMX-CD4 and the TR expression vector pMD18-T TR were from Addgene (Watertown, MA) and Sino Biological Inc. (Wayne, PA), respectively. Anti-FLAG, anti-HA, and anti-tubulin antibodies, and anti-HA- and anti-FLAG-conjugated agarose beads were purchased from Sigma (St. Louis, MO). The following siRNAs targeting human genes were obtained: ATPV0C from Sant Cruz Biotechnology (Dallas, Texas), ATP6V0C” from OriGene Technologies Inc., tetherin from Qiagen (Germantown, MD), and non-targeting control-siRNA from Dharmacon Inc. (Chicago, IL, now part of Horizon Discovery). Anti-human TR antibody was purchased from Zymed Laboratories Inc. (San Francisco, CA), anti-CD4 and anti-CD63 antibodies were from Santa Cruz, and anti-LAMP-1 from BD Biosciences (San Jose, CA). Bafilomycin was obtained from Tocris Bioscience (Minneapolis, MN). Zenon antibody labeling kits and the Alexa Fluor 488, 594, and 647 conjugated secondary antibodies were from Invitrogen (Grand Island, NY). Anti-Vpu (34), anti-BST2 (101), and anti-HIV-1 Ig were obtained from the NIH AIDS Reagent Program.

### Yeast two-hybrid screening

The yeast two-hybrid screen was carried out at Myriad Genetics (Salt Lake City, UT) as described previously (48). Briefly, the PNY200 yeast strain expressing the Vpu-fused GAL4 DNA binding protein was mated with the BK100 yeast strain transformed with a prey cDNA library (isolated from brain, spleen, and macrophages). After mating, the cells were plated onto selective media and the positive colonies were picked from the selection plates (102). The prey plasmids were isolated from the positive colonies, and the prey inserts were identified by sequence analysis. The interactions were confirmed by transforming the bait and prey plasmids into naïve yeast cells and monitoring the interactions by the chemiluminescent reporter gene assay system as described previously (102).

### Cell culture and transfection

293T and HeLa cell lines were maintained in Dulbecco-modified Eagle’s medium (DMEM) containing 10% or 5% fetal bovine serum (FBS), respectively. One day after plating, cells were transfected with appropriate plasmid DNA using Lipofectamine 2000 (Invitrogen Corp., Carlsbad, CA) according to the manufacturer’s recommendations. Twenty-four h posttransfection, virions were pelleted in an ultracentrifuge and cell and virus pellets were lysed and used for further analysis. In bafilomycin treatment experiments, 8 h posttransfection, cells were either untreated or treated with bafilomycin for 18 h prior to cell lysis.

### Immunoprecipitation and western blotting

For co-immunoprecipitation assays, 293T cells were transfected with FLAG-tagged ATP6V0C in the absence and presence of vectors expressing Vpu and human tetherin. After 24 to 30 h post-transfection, cells were lysed in 0.5% IGEPAL and immunoprecipitations were performed with agarose beads conjugated with anti-FLAG, anti-Vpu, or anti-HA antibodies. After overnight incubation, complexes were washed with 0.1% IGEPAL, and both cell lysates and immunoprecipitated samples were subjected to immunoblotting with anti-FLAG, anti-Vpu, and anti-HA antibodies. For immunoblot analyses, cell and virus pellets were lysed in a buffer containing 50 mM Tris-HCl (pH 7.4), 150 mM NaCl, 1 mM EDTA, 0.5% Triton X-100, and protease inhibitor cocktail (Roche life sciences, Basel, Switzerland). Proteins were denatured by boiling in sample buffer and subjected to SDS-PAGE, transferred to PVDF membrane, blocked with 5% milk and incubated with appropriate antibodies as described in the text. Membranes were then incubated with HRP-conjugated secondary antibodies, and chemiluminescence signal was detected by using West Pico or West Femto Chemiluminescence Reagent (Thermo Scientific, Waltham, MA). The protein bands were quantified by using Imagelab-Chemidoc (Bio-rad Laboratories, France).

### Virus release assays

One day after plating, 293T cells were transfected with WT or Vpu-defective pNL4-3 molecular clones in the absence and presence of ATP6V0C and tetherin expression vectors. One-day posttransfection, virions were pelleted in an ultracentrifuge and cell and virus pellets were lysed (103). To knock down ATP6V0C in HeLa and 293T cells, and tetherin in HeLa cells, one day after plating, cells were transfected with 50 nM non-target or ATP6V0C or tetherin siRNA with Oligofectamine transfection reagent (Invitrogen) in serum-free DMEM. After 6 to 7 h, DMEM containing 15% FBS was added and cells were cultured overnight. The next day, cells were infected with VSVG-pseudotyped wild-type (WT) or Vpu-defective HIV-1; after 8 h the medium was changed, and the siRNA transfection was repeated. Two days later, virions were pelleted in an ultracentrifuge and cell and viral pellets were lysed as above. Viral proteins in cell and virus lysates were immunoblotted with HIV-Ig (48) and virus release efficiency was calculated as the amount of virion-associated p24 as a fraction of total (cell-associated p24 and Pr55 plus virion-associated p24) Gag.

### Pulse-chase analysis

293T cells were transfected with HA-tagged human tetherin in the absence and presence of ATP6V0C expression vector. One day posttransfection, cells were pulse-labeled with [^35^S]Met-Cys for 30 min and washed with DMEM containing 10% FBS and removed from the dish in the same medium. Cells were split into five equal parts and incubated for 0, 0.5, 1, 2, 4 h. Cells were spun down after the indicated incubation time and lysed in 0.5% IGEPAL-containing lysis buffer. The cell lysates were subjected to immunoprecipitation with anti-HA antibody-tagged beads overnight, washed and analyzed by SDS-PAGE followed by fluorography (104).

### Immunofluorescence microscopy

For microscopy studies, 293T cells were cultured in chamber slides. One-day after plating, cells were transfected with human tetherin expression vector in the absence or presence of ATP6V0C expression vector. After 24 h, cells were rinsed with PBS and fixed with 3.7% paraformaldehyde in PBS for 30 min. The cells were rinsed with PBS three times, permeabilized with methanol at 20°C for 4 min, washed in PBS and incubated with 0.1 M glycine-PBS for 10 min to quench the remaining aldehyde residues. After blocking with 3% BSA-PBS for 30 min, cells were incubated with anti-HA and anti-FLAG antibodies diluted in 3% BSA-PBS for 1 h. The cells were washed with PBS three times and then incubated with secondary antibody conjugated with Alexa Fluor 488 and Alexa Fluor 594 diluted in 3% BSA-PBS for 1h to label anti-HA and anti-FLAG, respectively. In triple antibody staining experiments, cells were first incubated with anti-FLAG antibody diluted in 3% BSA-PBS for 1 h, then cells were washed with PBS three times and then incubated with secondary antibody conjugated with Alexa Fluor 647 diluted in 3% BSA-PBS. After washing with PBS three times, cells were incubated with Zenon Alexa Fluor 488-labeled anti-HA and Zenon Alexa Fluor 594-labeled anti-CD63 or anti-LAMP-1 for 1h. Finally, after washing with PBS three times, cells were mounted with Vectashield mounting media with DAPI (Vector Laboratories) and examined with a Delta-Vision RT microscope.

## Acknowledgements

We thank Myriad Genetics (Salt Lake City, UT) for carrying out yeast two-hybrid screening. We thank members of the Freed laboratory for helpful discussion and critical review of the manuscript. We thank K. Strebel and P. Bieniasz for their generous gifts of plasmids. The HIV-1 Ig, anti-Vpu and anti-human tetherin antisera were obtained from the NIH AIDS Reagent Program. Research in the Freed laboratory is supported by the Intramural Research Program of the Center for Cancer Research, National Cancer Institute, NIH.

## Author contributions

A.W. planned, designed and performed the experiments. M.S., A.K., A.G., and A.M. helped in biochemical analyses. E.O.F. coordinated and supervised the project. A.W. and E.O.F. wrote the manuscript.

## Conflict of interest

The authors declare no conflict of interest.

---

**Fig. S1.**
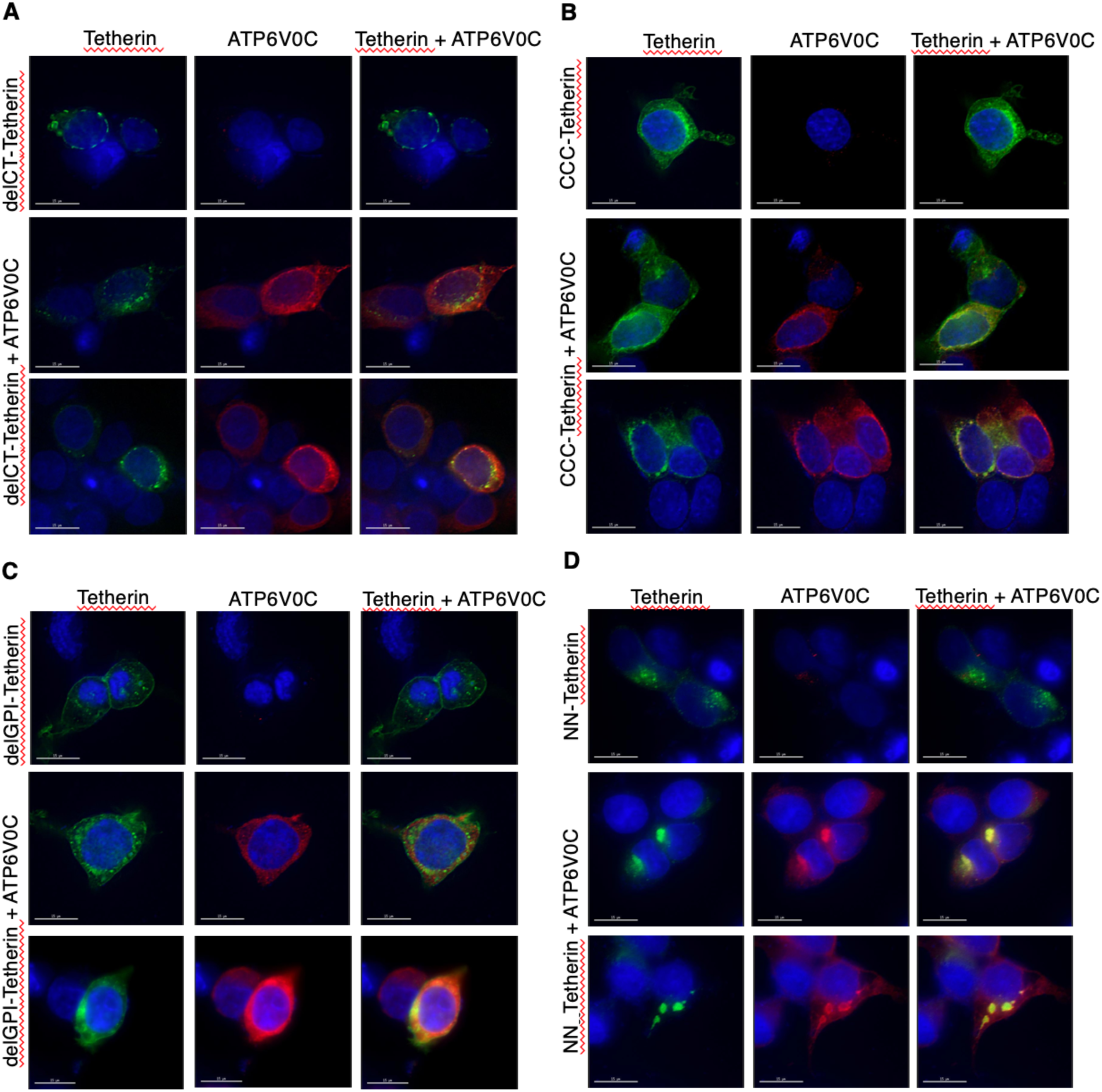
Localization of tetherin mutants in the presence of ATP6V0C. 293T cells were transfected with HA-tagged mutant tetherin expression vector alone or in combination with the FLAG-tagged ATP6V0C expression vector. Tetherin mutants tested include: (**S1A**) a cytoplasmic tail deletion mutant (delCT); (**S1B**) a non-dimerizing mutant (CCC); (**S1C**) a GPI-anchor deletion mutant (delGPI); and (**S1D**) a non-glycosylated mutant (NN). Twenty-four h post-transfection cells were fixed, stained as in Fig. 8A and images were acquired with a Delta-Vision RT deconvolution microscope. Scale bars, 15 μm.

